# Dual-responsive synthetic gene circuit for dynamic biologic drug delivery via inflammatory and circadian signaling pathways

**DOI:** 10.1101/2025.03.20.644403

**Authors:** Amanda Cimino, Fiona Pat, Omolabake Oyebamiji, Christine T.N. Pham, Erik D. Herzog, Farshid Guilak

**Author notes:** Corresponding Author: Farshid Guilak **Email:**.

## Abstract

**Background:** Engineered cells provide versatile tools for precise, tunable drug delivery, especially when synthetic stimulus-responsive gene circuits are incorporated. In many complex disease conditions, endogenous pathologic signals such as inflammation can vary dynamically over different time scales. For example, in autoimmune conditions such as rheumatoid arthritis or juvenile idiopathic arthritis, local (joint) and systemic inflammatory signals fluctuate daily, peaking in the early morning, but can also persist over long periods of time, triggering flare-ups that can last weeks to months. However, treatment with disease-modifying anti-rheumatic drugs is typically provided at continuous high doses, regardless of disease activity and without consideration for levels of inflammatory signals. In previous studies, we have developed cell-based drug delivery systems that can automatically address the different scales of flares using either chronogenetic circuits (i.e., clock gene-responsive elements) that can be tuned for optimal drug delivery to dampen circadian variations in inflammatory levels or inflammation-responsive circuits (i.e., NF-κB-sensitive elements) that can respond to sustained arthritis flares on demand with proportional synthesis of drug. The goal of this study was to develop a novel dual-responsive synthetic gene circuit that responds to both circadian and inflammatory inputs using OR-gate logic for both daily timed therapeutic output and enhanced therapeutic output during chronic inflammatory conditions.

**Results:** We developed a synthetic gene circuit driven by tandem inflammatory NF-κB and circadian E’-box response elements. When engineered into induced pluripotent stem cells that were chondrogenically differentiated, the gene circuit demonstrated basal-level circadian output with enhanced stimulus-responsive output during an inflammatory challenge shown by bioluminescence monitoring. Similarly, this system exhibited enhanced therapeutic levels of biologic drug interleukin-1 receptor antagonist (IL-1Ra) during an inflammatory challenge in differentiated cartilage pellets. This dual-responsive therapeutic gene circuit mitigated both the inflammatory response as measured by bioluminescence reporter output and tissue-level degradation during conditions mimicking an arthritic flare.

**Conclusions:** The dual-responsive synthetic gene circuit developed herein responds to input cues from two key homeostatic transcriptional networks, enabling dynamic and tunable output. This proof-of-concept approach has the potential to match drug delivery to disease activity for optimal outcomes that addresses the complex environment of inflammatory arthritis.

## Introduction

Engineered cell-based systems can act as living devices that sense and respond to their environments to achieve dynamic, personalized treatment. Using synthetic biology tools, cellular machinery can be repurposed to design biologic-based circuitry analogous to electrical circuitry. Synthetic gene circuits have been previously used to generate anti-inflammatory therapeutic drugs based on input cues including inflammatory activation, mechanical load (“mechanogenetic”), or circadian time of day (“chronogenetic”)^1-4^. By harnessing endogenous signaling pathways responsive to these cues, gene circuits consisting of a promoter that transcriptionally responds to the selected cue and a downstream gene encoding a biologic can be genetically edited to a cell’s DNA. Subsequently, the introduced gene circuit enables the cell to sense its environment and respond therapeutically. Since these circuits function based on endogenous cellular pathways, they are inherently able to tune therapeutic output and incorporate aspects like feedback in response to the dynamic environment. Additionally, increasingly complex gene circuits have been designed to incorporate elements like genetic amplifiers, switches, and Boolean logic operations to address multifaceted disease scenarios ^5, 6^.

Rheumatoid arthritis (RA) and juvenile idiopathic arthritis (JIA) are debilitating autoimmune inflammatory diseases that effect joints throughout the body, causing painful swelling, cartilage degradation, and bone erosion ^7-10^. Moreover, the severity of inflammation can change over time, making it particularly challenging to properly dose therapeutic drugs. On a short-term scale, systemic inflammatory mediators and symptoms like joint stiffness cycle in daily rhythms, peaking in the early morning ^11^. On a longer-term scale, inflammatory arthritis patients experience unpredictable “flare-ups,” lasting periods of weeks to months during which overall symptoms significantly worsen ^10, 12^. Currently, disease modifying antirheumatic drugs (DMARDs), the standard of care for RA, are prescribed at high, immunosuppressive doses regardless of dynamic inflammation ^10, 13, 14^. Consequently, the incongruence in treatment intensity and disease tempo leads to suboptimal arthritis control while contributing to potential adverse effects such as heightened risk of infection ^15-17^.

While protein delivery systems have many advantages for controlled drug delivery, current approaches do not have adequate capacity for dynamic and individualized stimulus-response or timed release ^18-21^. To address the different time scales of RA flares, chronogenetic circuits can be tuned for optimal daily delivery to dampen the early-morning rise in inflammation, while inflammatory-responsive circuits can target unpredictable sustained flares proportionally with dynamic negative feedback (**Fig. 1**) ^2, 3, 22, 23^. However, these individual strategies may not meet the needs of RA therapeutic delivery across time scales. For example, inflammatory-responsive gene circuits cannot preemptively generate therapeutics to dampen rising circadian inflammation prior to sensing it, and chronogenetic circuits depend on robust transcriptional-translational circadian cycles, which are disrupted during highly inflammatory conditions, potentially contributing to the loss of circadian regulation in RA joints ^24-26^. Therefore, the overall goal of this work was to develop a dual-responsive synthetic gene circuit that responded to both circadian and inflammatory cues using OR-gate logic for daily timed therapeutic output with enhanced therapeutic output during inflammatory conditions. This was accomplished by creating a gene circuit promoter that contained both NF-κB inflammatory response elements and E’-box circadian elements. For dynamic, feedback controlled inflammatory response, NF-κB transcription factors bind their response elements to induce transcriptional activation during inflammatory conditions ^2^. To generate daily prescribed release, BMAL1 and CLOCK proteins bind E’-boxes to activate clock-controlled genes, which include PER and CRY proteins that inhibit BMAL1/CLOCK activation, leading to an autoregulated 24-hour cycling of activated gene expression at this element ^22^. Together, this system results in a dual-responsive circuit for biologic production using OR-gate logic.

**Figure 1.**
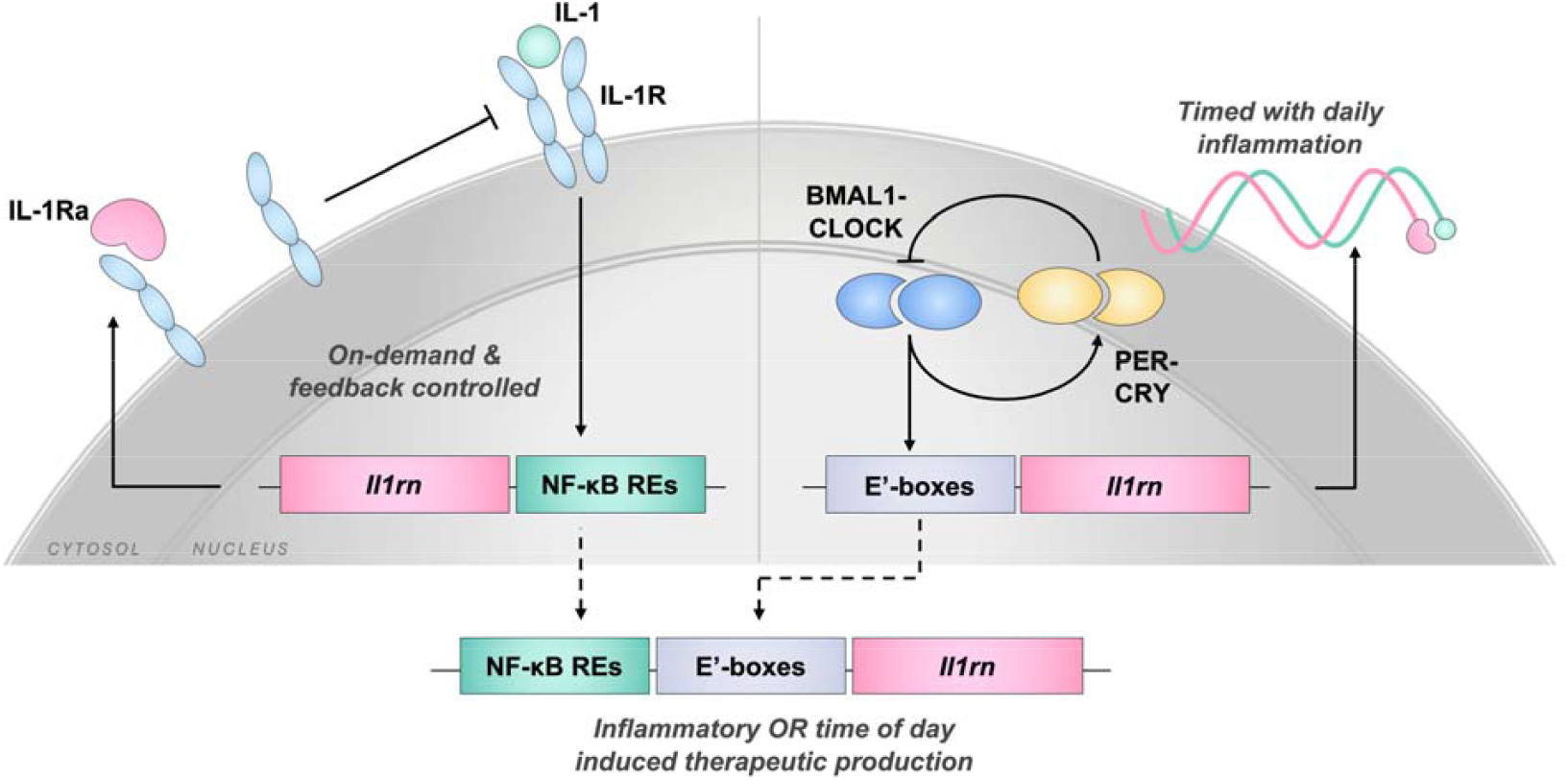
The inflammatory and circadian dual-responsive gene circuit produces a biologic using OR-gate logic. Inflammatory-driven biologic production is programmed by NF-κB response elements (REs) that promote downstream gene activation during inflammatory signaling, such as when IL-1 binds its receptor (IL-1R). This leads to IL-1Ra production that blocks inflammatory signaling, providing on-demand, feedback controlled therapeutic delivery. Chronogenetic delivery is driven by the central internal circadian feedback loop, where binding of BMAL1-CLOCK dimers in E’-box elements activates downstream gene expression, as well as the PER and CRY proteins that dimerize to block BMAL1-CLOCK activation, generating a self-regulated timing mechanism. Using E’-box elements upstream of the therapeutic gene permits times daily delivery that can coincide with daily changes in inflammation. Combing these approaches, the dual-input circuit utilizes NF-κB REs followed by E’-boxes to drive therapeutic gene expression that responses to inflammatory signaling or programmed time of day.

## Materials and Methods

### Gene circuit design

Inflammatory-responsive (NFκB-LUC, NFκB-IL1Ra) and circadian-responsive (E’box-GFP-LUC, E’box-IL1Ra) circuits were designed as previously described ^2, 22^. Briefly, the NF-κB-driven circuits consisted of a promoter with five canonical NF-κB recognition motifs derived from promoters of inflammatory-responsive genes (*Infb1, Il6, Mcp1, Adamts5*, and *Cxcl10*), an upstream negative regulatory element to repress background expression, and a downstream minimal CMV. The E’-box-driven circuits had promoters composed of three tandem E’-boxes derived from the *Per2* promoter followed by a minimal CMV. To construct the dual-responsive inflammatory-circadian circuit with an NFκB.E’box promoter, the five NF-κB recognition motifs were isolated by PCR amplification from NFκB-LUC with the addition of homology arms to match the targeted insert region directly upstream of the three tandem E’-boxes in E’box-GFP-LUC or E’box-IL1Ra (*5’-TCACATAGTGGAAAACGTGACCGCGCGCGCACTAGGATCTGGGAAGTCCCCTCGA, 5’-ATTACAAAAACAAATTACAAAATTCAAAATTTTATCGATAAGAGGTACCGAGCTCTT ACG*). E’box-GFP-LUC and E’box-IL1Ra were digested with SpeI to open the vector at the insert site. Then, NFκB.E’box-GFP-LUC and NFκB.E’box-IL1Ra were cloned by Gibson Assembly. Correct insertion was confirmed by Sanger sequencing.

### Induced pluripotent stem cell (iPSC)-derived cartilage differentiation and culture

Cartilage pellets were differentiated according to a previous developed protocol ^27^. First, iPSCs were cultured on mitomycin C-treated mouse embryonic fibroblasts (MEFs) for five days in gelatin-coated dishes in culture media consisting of Dulbecco’s Modified Eagle Medium-high glucose (DMEM-HG), 20% lot-selected serum, 1% minimum essential medium (MEM) non-essential amino acids, 55 μM 2-Mercaptoethanol, 25 μg/mL gentamicin, and 1,000 units/mL mouse leukemia inhibitory factor (LIF). MEFs were removed by feeder cell subtraction, and then cells were plated as high-density micromass cultures for 14 days in chondrogenic media (DMEM-HG; 1% insulin, transferrin and selenous acid+ (ITS+); 1% MEM non-essential amino acids; 1% penicillin/streptomycin; 55 μM 2-Mercaptoethanol; 50 μg/mL ascorbate, and 40 μg/mL proline) supplemented with 50 ng/mL bone morphogenic protein 4 (BMP-4) and 100 nM dexamethasone on days three and five of micromass culture. Micromasses were digested with pronase and collagenase II and plated onto gelatin-coated flasks as pre-differentiated iPSCs (PDiPSCs). PDiPSCs were expanded (DMEM-HG, 10% lot-selected serum, 1% ITS+, 1% MEM non-essential amino acids, 1% penicillin/streptomycin, 55 μM 2-Mercaptoethanol, 50 μg/mL ascorbate, 40 μg/mL proline, and 4 ng/mL basic fibroblast growth factor), collected, and pelleted in 15-mL conical tubes at 250,000 cells per pellet culture. Pellets were maintained for 21 days in chondrogenic media supplemented with 100 nM dexamethasone, 50 μg/mL ascorbate, 40 μg/mL proline, and 10 ng/mL transforming growth factor-β3 (TGF-β3).

Prior to testing, cells were synchronized in media with 100 nM dexamethasone for one hour and then cultured in media without dexamethasone or growth factors to reduce potential impacts on synchronization or inflammatory response. Inflammatory challenges were induced by adding 0.1 or 1 ng/mL IL-1β to the culture media after synchronization.

### Lentivirus production and cell transduction

Second-generation packaged vesicular stomatitis virus glycoprotein pseudotyped lentivirus was produced according to a standard protocol ^28^. HEK293T cells were transfected by calcium phosphate precipitation with the psPAX2 packaging vector, pMD2.G envelope protein vector, and expression vectors. HEK293T culture media was collected, filtered, and stored at −80°C until use. A viral titer was performed in HeLa cells to identify the multiplicity of infection (MOI). Chondrogenic cells were transduced at the PDiPSC stage for 24 hours at an MOI of approximately 3 in media supplemented with 4 μg/mL polybrene.

### Bioluminescence circuit characterization

Bioluminescence was recorded at 15-minute intervals as relative luminescence units (RLU) in a light-protected enclosed CO_2_-controlled incubator. After synchronization, cells were cultured in phenol-free media supplemented with 100 μM luciferin for recordings. During inflammatory challenge, recording media was replaced with fresh media containing 0, 0.1, or 1 ng/mL IL-1β. When recording pellets, the first two hours after a media change were discarded due to the strong initial peak from adding fresh luciferase. Area under the curve (AUC) was quantified using Prism GraphPad. Relative peak value post-challenge was determined by the maximum value for each sample. Normalization was performed with respect to the non-challenged control for each group. For circadian signals, a sinusoidal curve was fit by Prism GraphPad (nonlinear fit for a sine wave with a nonzero baseline, least squares regression, constrained to wavelength <18) to determine the fit curve’s period and baseline over at least 48 hours.

### Gene expression

Samples were frozen at −80°C until RNA isolation. Pellets were homogenized using a miniature bead beater, and RNA was then isolated according to the manufacture’s protocol (Norgen Biotek). Complementary DNA (cDNA) was produced using Invitrogen SuperScript VILO cDNA master mix. Gene expression was determined by quantitative reverse transcription polymerase chain reaction (RT-qPCR) using Applied Biosystems Fast SyBR Green master mix according to the manufacture’s protocol. The ΔΔCT method was used to determine relative fold change in gene expression, with respect to endogenous expression of *Gapdh*. Primers for were synthesized by Integrated DNA Technologies and verified for efficiency (Table 1).

**Table 1.**
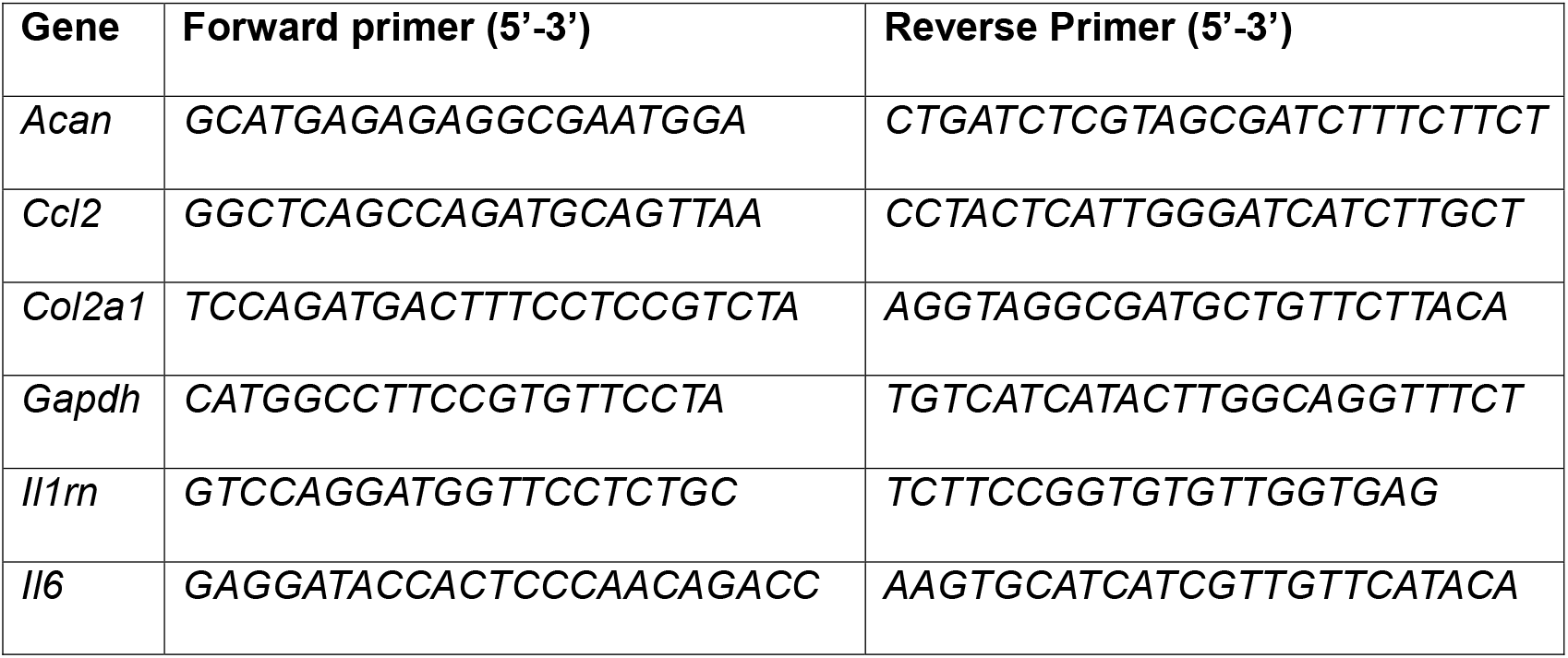
Primers for RT-qPCR expression.

### Enzyme-linked immunosorbent assay

Culture media was collected and stored at −20°C until analysis. Concentrations of IL-1Ra in the media were quantified in duplicate using the R&D Systems DuoSet enzyme-linked immunosorbent assay (ELISA) for mouse IL-1Ra with absorbance readings at 450 nm and 540 nm.

### Biochemical analysis of pellets

Samples were collected and stored at −20°C until digestion. Pellets were digested in papain overnight at 65°C. Total sulfated glycosaminoglycan (GAG) content in the pellets was quantified using a 1,9-dimethylmethylene blue (DMMB) assay and normalized to total DNA content, quantified by a PicoGreen assay (Thermo Fisher) according to the manufacturer’s protocol.

### Histological analysis of pellets

To prepare histological samples, pellets were fixed in a formalin solution, dehydrated using ethanol, and embedded in paraffin. 8-μm thick sections were stained with Safranin-O, Fast-Green, and hematoxylin. Brightfield images of slides were taken at 20x magnification using an Olympus VS120 microscope.

### Statistical analysis

Metrics quantified from bioluminescent signals (area under the curve (AUC), baseline, period) were assessed by one-way ANOVA with Tukey’s multiple comparison test or t-test. Time course data (*Il1rn*, IL-1Ra) was assessed by two-way ANOVA with Sidak’s multiple comparison test between timepoints. Protein accumulation, proteoglycan content, and gene expression at 24-or 72-hours post-challenge were assessed by one-way ANOVA with Tukey’s multiple comparison test.

## Results

### Dual NF-κB- and E’-box-responsive promoter generates basal-level circadian output with on-demand enhanced output during inflammatory challenge

Bioluminescent reporter systems were investigated to characterize the dynamic expression kinetics of the gene circuits in the presence or absence of inflammatory challenges that mimic arthritic flares *in vitro* in monolayer PDiPSCs. The NF-κB-driven system responded on demand to inflammatory challenge, whereas the E’-box-driven system autonomously cycled on and off due to circadian regulation (**Fig. 2A**). As shown by bioluminescent reporters, NF-κB-driven output (NFκB-LUC) increased with challenge, demonstrated by relative AUC that was proportional to the concentration of IL-1β (**Fig. 2B, C**), without any observed circadian expression trends. Inversely, E’-box-driven chronogenetic output (E’box-GFP-LUC) was suppressed by IL-1β challenge, shown by reduced AUC and baseline of the sinusoidal curve fit and increased period length (**Fig. 2D-G**).

**Figure 2.**
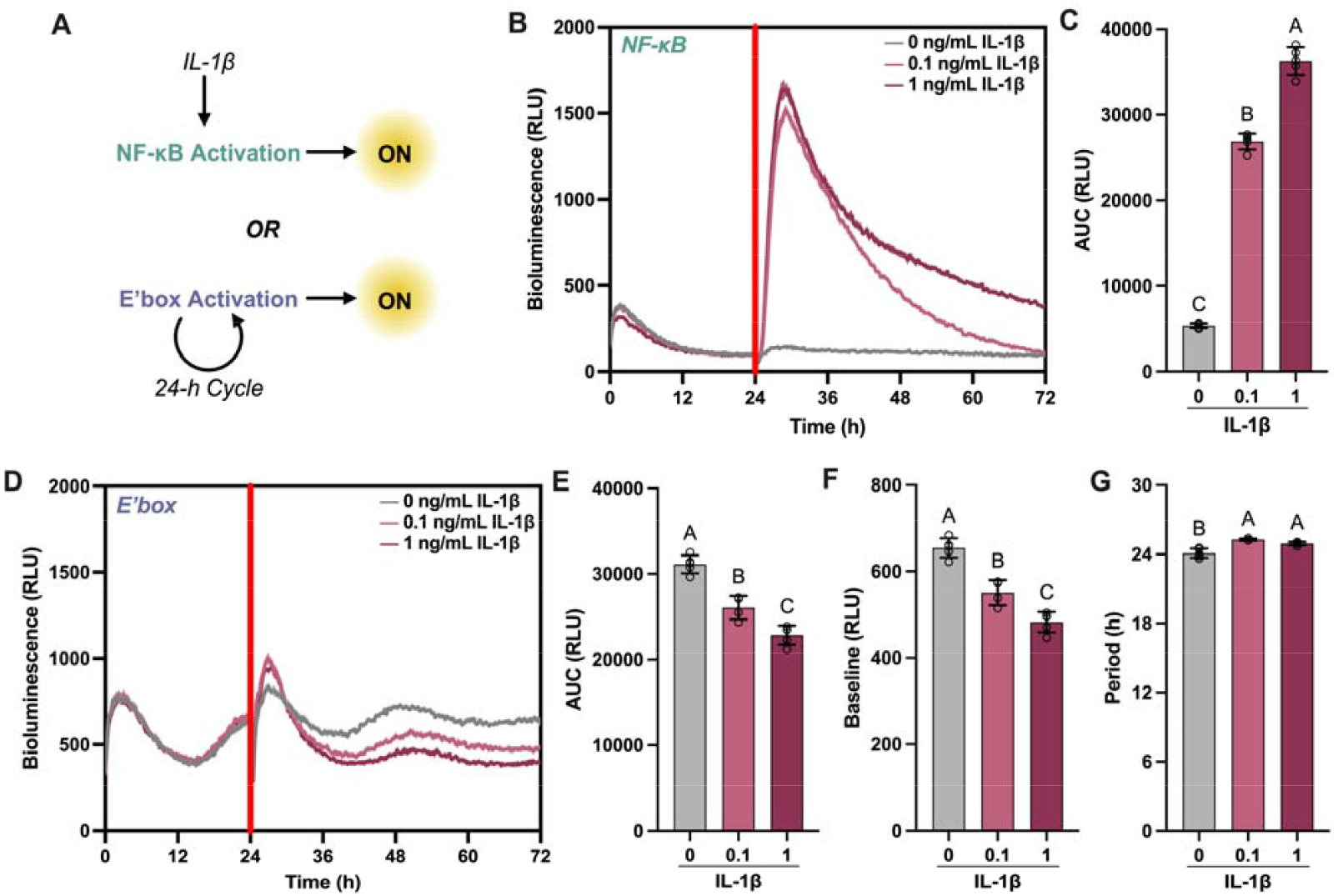
Distinct, disease-relevant cues activate inflammatory or chronogenetic gene circuits. (A) The inflammatory-driven NF-κB circuit activates on-demand in response to IL-1β challenge, whereas the circadian E’-box circuit is autonomously regulated by clock-controlled feedback on a 24-hour on/off cycle. (B) Dynamic tracking of gene circuit response after inflammatory challenge (indicated by the red line at 24 hours) showed an increased response for NFκB-LUC, (C) quantitatively demonstrated by increased area under the curve (AUC). (D) Chronogenetic output by E’box-GFP-LUC was dampened with IL-1β challenge, demonstrated by reduced (E) AUC and (F) baseline of the sinusoidal curve fit and (G) lengthened period. Figures show mean and SEM, n=4-5/condition. Groups not sharing the same letter are significant (p<0.05) by one-way ANOVA with Tukey’s multiple comparisons test.

To combine these approaches and overcome individual limitations, the dual-responsive circuit (NFκB.E’box-GFP-LUC) was constructed and showed basal-level circadian output in the absence of an inflammatory challenge, shown by a 24-hour period, but had enhanced output proportional to the challenge, quantified by AUC, due to it responsive to inflammatory challenge (**Fig. 3A-C**). Consequently, the dual-responsive NF-κB- and E’-box-driven promoter offers a combined mode of activation for therapeutic delivery during RA flares.

**Figure 3.**
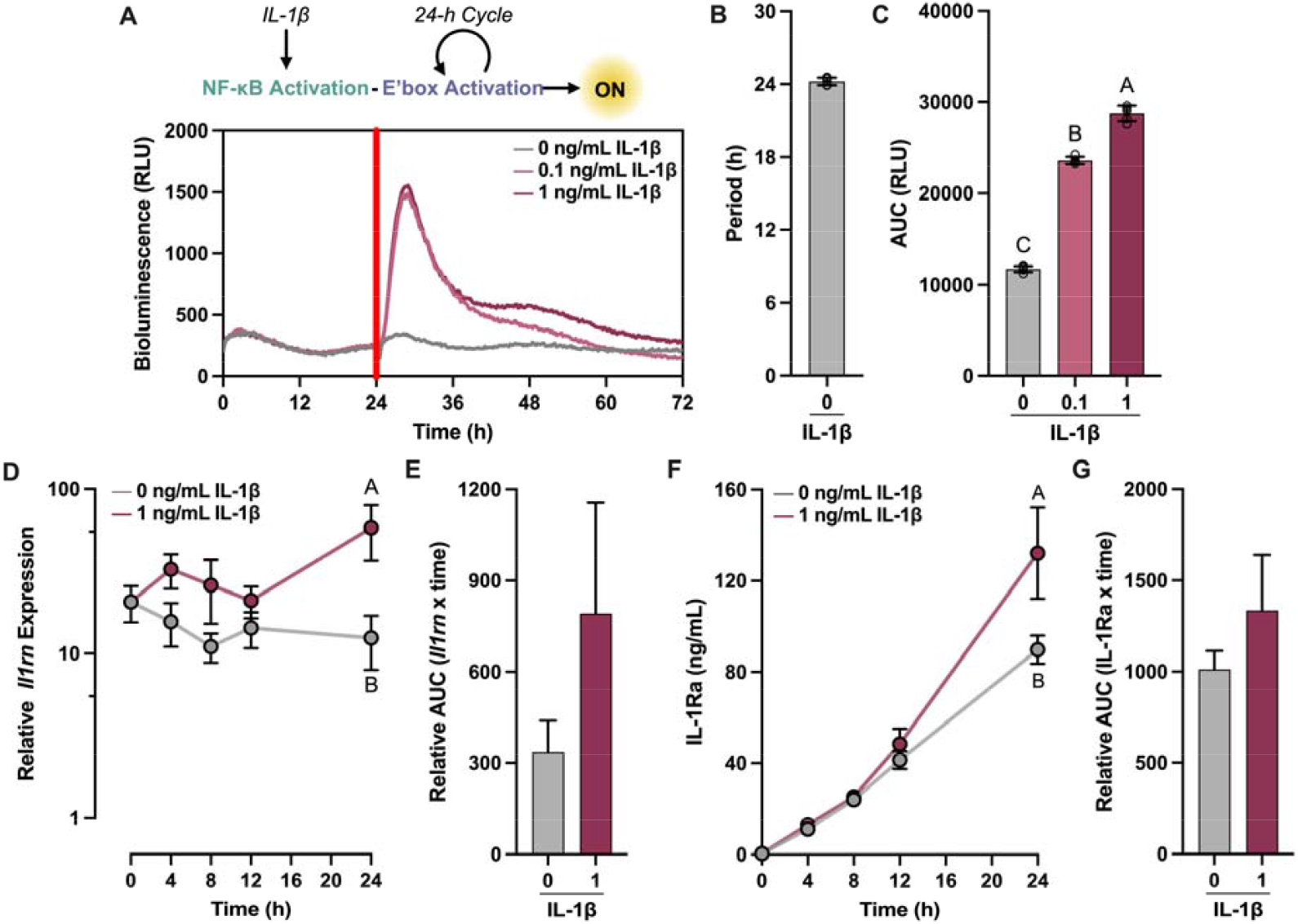
The dual-responsive inflammatory and chronogenetic design activates in response to either cue. (A) Continuous bioluminescence monitoring of NFκB.E’box-GFP-LUC response reveals basal-level circadian output prior to challenge and in the absence of stimulation, shown by the signal’s (B) 24-hour period. (C) When challenged, circuit activation increased, demonstrated by AUC. In alignment with bioluminescence, (D) *Il1rn* expression and (F) IL-1Ra accumulation was increased with challenge in comparison to non-challenged controls, resulting a significant difference between groups by 24 hours post-challenge and (E,G) increased AUC. Figures show mean and SEM, n=3-7/condition. Different letters indicate significant differences (p<0.05) by one-way ANOVA with Tukey’s multiple comparisons test (B, C) or two-way ANOVA with Sidak’s multiple comparisons test (D, F).

### NFκB.E’box-IL1Ra generates enhanced therapeutic output in response to IL-1β challenge

To assess the potential therapeutic utility of the circuit, the expression profile of IL-1Ra was evaluated in differentiated cartilage pellets with and without challenge. Previous work has shown that E’box-IL1Ra generated IL-1Ra on a circadian basis with a rate of change that fit a 24-hour period ^22^. Likewise, NFκB-IL1Ra produced IL-1Ra proportional to the level of IL-1 challenge ^2^. Since the dual-responsive designs generates output due to both circadian and inflammatory cues, we determined if there was a differential response in the presence or absence of an inflammatory challenge during the first 24 hours of activation that capture when both cues will likely be present, based on bioluminescence data and previous studies ^2, 22^. As expected, IL-1Ra concentration in the culture media increased over time in the presence or absence of a challenge and reached similar a magnitude as circadian E’-box-driven circuits ^22^. However, by 24 hours post-challenge, the challenged group had significantly increased *Il1rn* expression and IL-1Ra production than the unchallenged group, indicative of enhanced activation due to NF-κB responsiveness (**Fig. 3D,F**). Likewise, this corresponded to a trend in increased relative AUC during challenge (**Fig. 3E,G**), representative of an approximately 30% greater expression or output of the therapeutic over the first 24 hours post-challenge.

### Dual-responsive therapeutic design dampens inflammatory activation

After confirming that the circuit responded to inflammatory activation with enhanced biologic production, we investigated the capacity of the therapeutic circuit to impact the inflammatory state of cartilage pellets. Using NFκB.E’box-GFP-LUC as a reporter for overall inflammation, we compared the level of inflammatory activation, quantified by AUC, in pellets transduced with only the reporter or with both the reporter and therapeutic versions of the circuit (**Fig. 4A, C**). Prior to challenge, there were no significant differences in AUC between groups. However, following challenge with 1 ng/mL IL-1β, there was a significant increase in inflammatory activation that was approximately two-or four-times greater with respect to the unchallenged controls in those with or without the therapeutic circuit, respectively (**Fig. 4B, D**). While this represented an increase over the respective controls, those with the therapeutic circuit had a significantly reduced AUC response when compared to groups without NFκB.E’box-IL1Ra (**Fig. 4E**). Additionally, the relative peak in bioluminescence post-challenge was significantly lower with the therapeutic circuit, achieving near baseline levels (**Fig. 4F**). Together, these findings demonstrate the capacity of NFκB.E’box-IL1Ra to mitigate the inflammatory activation.

**Figure 4.**
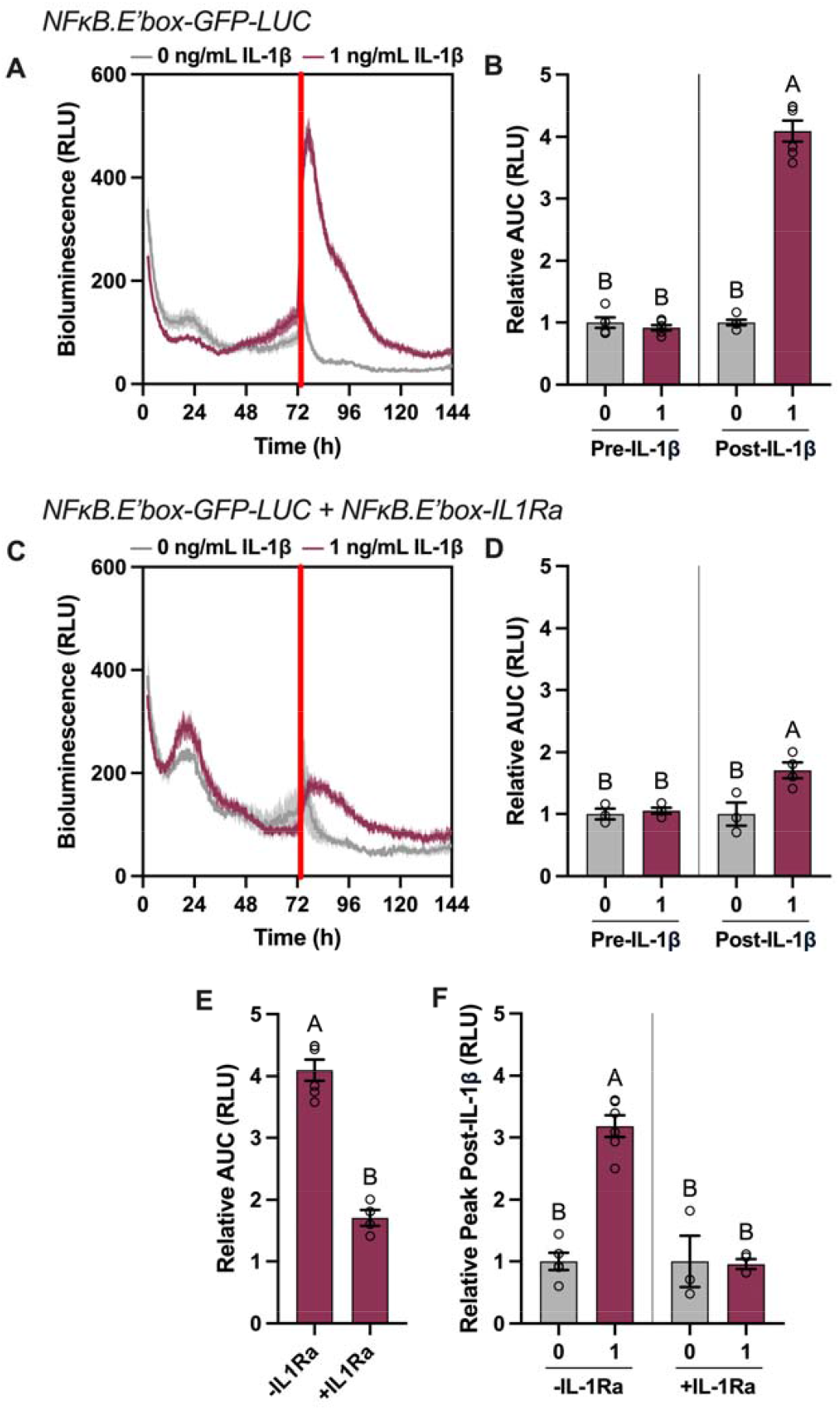
Therapeutic production from NF_κ_B.E’box-IL1Ra mitigated inflammatory activation response. Continuous bioluminescence monitoring of NFκB.E’box-GFP-LUC response (A) without or (C) with the inclusion of NFκB.E’box-IL1Ra demonstrated reduction of the circuit activation following challenge quantified by (B, D, E) AUC and (F) relative peak activation. Figures show mean and SEM, n=3-6/condition. Groups not sharing the same letter are significant (p<0.05) by one-way ANOVA with Tukey’s multiple comparisons test (B, D, F) or t-test (E).

### NFκB.E’box-IL1Ra protects an *in vitro* model of arthritis from inflammatory degradation

To assess an *in vitro* RA model, the therapeutic gene circuit was compared to non-transduced (NT) controls that represented the native tissue in cartilage pellets at 24- and 72-hours following challenge with IL-1β. The dual-responsive circuit demonstrated significantly more IL-1Ra output in the presence of a challenge in comparison to its unchallenged control at both time points, underscoring its capacity to enhance therapeutic delivery during inflammatory conditions (**Fig. 5A, B**). This aligned with protection of the tissue matrix, both quantitatively by sulfonated glycosaminoglycan (GAG) content that demonstrated near complete protection and qualitatively by maintained histological staining of proteoglycans by red Safranin-O (**Fig. 5C, D**). Additionally, the loss of cartilage gene expression and the activation of inflammatory gene expression were reduced in comparison to non-transduced controls (**Fig. 5E**). In summary, this data demonstrates that tissue-level protection is provided by the dual-responsive circuit design during conditions mimicking an arthritic flare.

**Figure 5.**
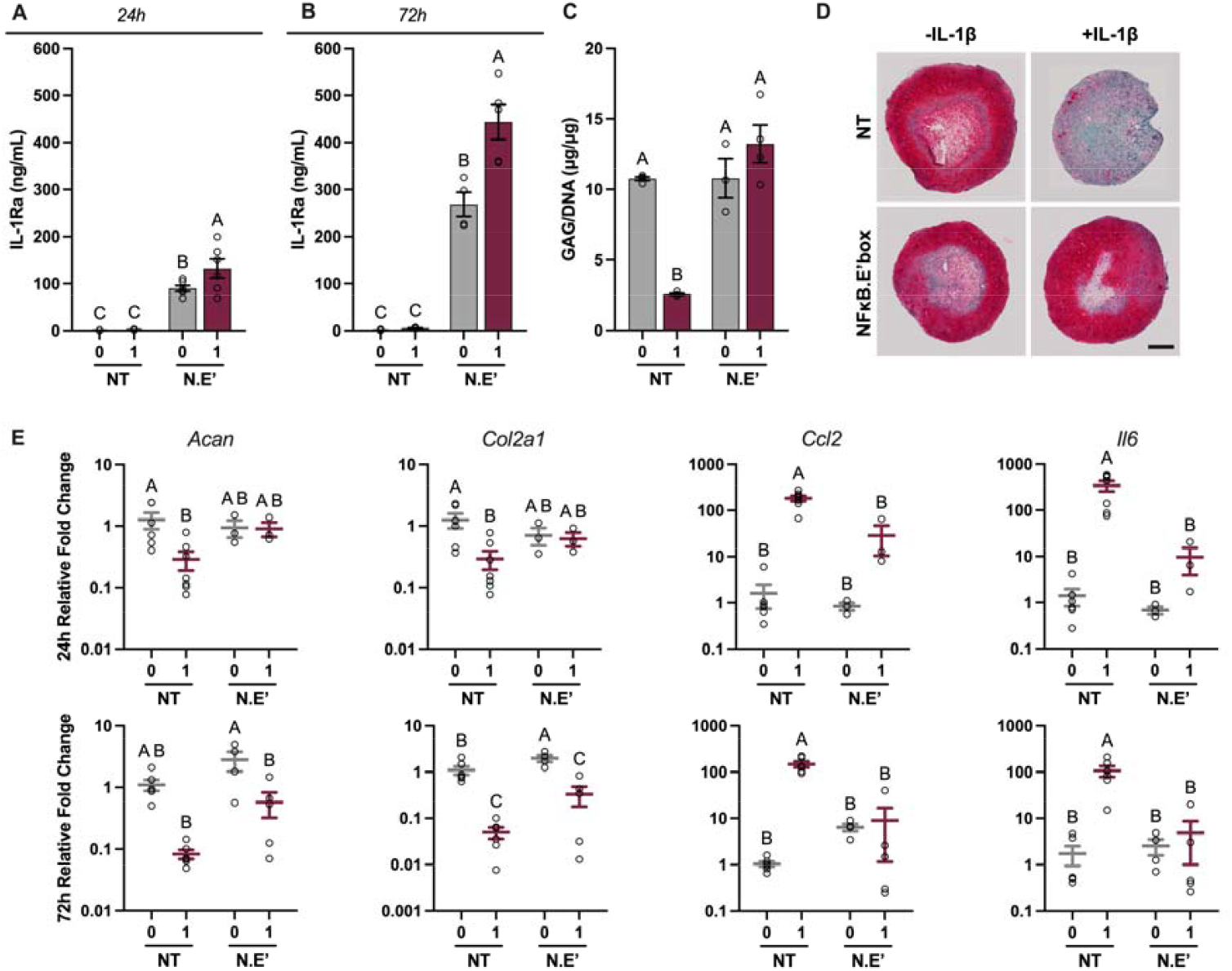
The dual-responsive gene circuit protected a model of arthritis from tissue-level damage. IL-1Ra production was significantly increased in comparison to non-transduced (NT) controls and with challenge at both (A) 24 hour and (B) 72 hours post-challenge. This contributed to protected cartilage matrix, shown by (C) quantified sulfonated GAG/DNA and (D) representative images of histological sections stained with Safranin-O, Fast-Green, and hematoxylin (n=2/condition; scale: 200μm). (E) At a gene expression level, loss of cartilage associated genes aggrecan (*Acan*) and type-II collagen (*Col2a1*) and increase in inflammatory associated genes C-C Motif Chemokine Ligand 2 (*Ccl2*) and interleukin-6 (*Il6*) were improved by the therapeutic circuit. Figures show mean and SEM, n=3-8/condition (A-C, E). Groups not sharing the same letter are significant by one-way ANOVA with Tukey’s multiple comparison test.

## Discussion

In this study, we designed and tested a novel synthetic gene circuit responsive to both circadian and inflammatory signaling pathways for both timed chronogenetic and inflammatory feedback-controlled cell-based drug delivery. This gene circuit successfully demonstrated both independent and dual-responsive synthesis of biologic drugs on demand. In the absence of an inflammatory challenge, this circuit responded to endogenous circadian transcriptional feedback signals through E’-box elements, resulting in autonomous cycling over a 24-hour period. During inflammatory challenge, this circuit had enhanced output that was regulated through a central inflammatory pathway, allowing the circuit to sense and respond to its dynamic environment. With this OR-gate digital logic, either inflammatory or time-of-day input resulted in disease-relevant activation for fine-tuned therapeutic release. Through this dual response, the circuit demonstrated a proof-of-concept design to address the complexity of flares in RA, JIA, or other inflammatory diseases that occur on both daily and long-term scales.

As shown by both bioluminescent reporters for real-time circuit activation kinetics and the therapeutic release profile of IL-1Ra at discrete time points, the NFκB.E’box promoter had significantly higher activation in the presence of an inflammatory challenge than in its absence. Without NF-κB responsiveness, we have previously shown that there was no significant change in IL-1Ra release during IL-1β challenge ^22^. However, this design permits enhanced delivery that can better address the dynamic long-term environment of inflammatory arthritis. Supporting the utility of this design, our *in vitro* model of arthritis was protected in both the level of inflammatory activation measured by bioluminescence reporters and the tissue-level matrix and gene expression in comparison to control samples.

Traditionally, therapeutic drugs are delivered at prescribed times, such as daily or weekly, without regard to the dynamic disease state, effectively functioning as open-loop systems ^29^. This approach contributes to a mismatch between the given and required dosage, which can lead to side effects while leaving the disease uncontrolled. In autoimmune conditions like RA and JIA, disease management can be exceptionally challenging, due to unpredictable flare ups in disease severity and medication responsiveness that can decrease over time ^12, 30-33^. With these challenges, closed-loop systems that use real-time input to provide therapeutic delivery when it is most beneficial may support improved disease control. Since cells have the necessary machinery to sense their environment and respond rapidly, adapting these tools to form gene circuits is a relatively straightforward extension of natural homeostatic mechanisms.

The goal of this study was to generate a system that provided dynamic control of biologic drug synthesis across multiple time scales that are characteristic of inflammation in various autoimmune conditions such as RA and JIA. With this design, the gene circuit can be introduced with a single promoter-output cassette. While existing iterations of gene circuits target distinct environmental cues, including inflammatory, circadian, or mechanical activation alone, the dual-responsive circuit addresses two interconnected disease-relevant conditions, achieving more tunable control ^1-3^. Others have successfully developed circuit systems in mammalian cells that include tools like amplification, feedback control, and digital logic for precise therapeutic delivery ^5, 6, 34^. These elements support more controlled delivery; however, to achieve this level of regulation, orthogonal components are often required, which increases system complexity. Herein, we developed a system that does not require the introduction of non-native signaling molecules or transcriptional machinery for circuit transduction. Instead, we demonstrated increased circuit control through a dual-responsive promoter that activates through endogenous signaling pathways, creating a simple but multifaceted system.

In summary, this dual-responsive synthetic gene circuit has the potential to address challenges in synthetic gene circuit control for inflammatory arthritis. Beyond arthritis, other inflammatory diseases are implicated to have associated inflammatory and circadian changes, including but not limited to inflammatory bowel diseases and chronic airway diseases ^35-37^. This proof-of-concept circuit design can be tailored to address a multitude of conditions. Herein, we demonstrated that E’-box-driven rhythms can be utilized in a dual-responsive system, but other phases in the 24-hour period can be targeted with circadian response elements like D-boxes or RREs ^22^. Moreover, combinations of these elements may be able to encode circadian expression at even more precise times ^38, 39^. Likewise, therapeutic output can be modulated based on the desired therapeutic protein, such as in previous efforts to generate circuits producing either IL-1Ra or soluble tumor necrosis factor receptor ^4^. Since this system was developed in a cartilage model, the design can be translated to *in vivo* studies, as cartilage is avascular and aneural, and can be implanted subcutaneously for long-term therapeutic delivery ^23, 40^. Therefore, this dual-responsive system serves as a prototype for future designs tailorable to specific disease scenarios.

## Conclusions

We developed a dual-responsive synthetic gene circuit for the delivery of an anti-inflammatory biologic that addresses both acute and prolonged changes in inflammatory signals. This was accomplished by repurposing elements of two main transcriptional networks that drive inflammatory and circadian gene expression, respectively. With this approach, the dual-responsive gene circuit generated a dynamic and tunable output, which has the potential to deliver drug in anticipation of daily changes in disease activity and in response to sustained flares for optimal outcomes.

## Declarations

### Ethics approval and consent to participate

Not applicable

### Consent for publication

Not applicable

### Availability of data and materials

All data generated or analyzed during this study are included in this published article.

### Authors’ contributions

AC and FG developed the concept of the study; AC, CP, EH, and FG designed the studies; AC, FP, and OO performed the studies and analyzed the data; AC and FG wrote the manuscript; all authors edited and approved the manuscript.

## Acknowledgements

Not applicable

